# Duplications of human longevity-associated genes across placental mammals

**DOI:** 10.1101/2023.05.02.539026

**Authors:** Zixia Huang, Chongyi Jiang, Jiayun Gu, Marek Uvizl, Sarahjane Power, Declan Douglas, Joanna Kacprzyk

## Abstract

Natural selection has shaped a wide range of lifespans across mammals, with a few long-lived species showing negligible signs of ageing. Approaches used to elucidate the genetic mechanisms underlying mammalian longevity usually involve phylogenetic selection tests on candidate genes, analyses of differential gene expression between age cohorts or species, and measuring age-related epigenetic changes. However, the link between gene duplication and evolution of mammalian longevity has not been widely investigated. Here, we explored the association between gene duplication and mammalian lifespan by analysing 287 human longevity-associated genes across 37 placental mammals. We estimated that the expansion rate of these genes is eight times higher than their contraction rate across these 37 species. Using phylogenetic approaches, we identified 43 genes whose duplication levels are significantly correlated with longevity quotients (FDR < 0.05). In particular, strong correlation observed for four genes (*CREBBP*, *PIK3R1*, *HELLS*, *FOXM1*) appears to be driven mainly by their high duplication levels in two ageing extremists, the naked mole rat (*Heterocephalus glaber*) and the greater mouse-eared bat (*Myotis myotis*). Further sequence and expression analyses suggest that the gene *PIK3R1* may have undergone a convergent duplication event, whereby the similar region of its coding sequence was independently duplicated multiple times in both of these long-lived species. Collectively, this study identified several candidate genes whose duplications may underlie the extreme longevity in mammals, and highlighted the potential role of gene duplication in the evolution of mammalian long lifespans.

## Significance

Long-lived mammals have naturally evolved exceptionally long lifespans and healthspans, which renders them as ideal unconventional models to ascertain the molecular basis of extended longevity. Despite an increase in our knowledge on mammalian longevity mechanisms, the link between gene duplication and evolution of mammalian longevity has not been widely explored. Here, we investigated the association between duplications of human longevity-associated genes and mammalian longevity across 37 mammals, potentially providing novel candidate genes for future functional validation of their biological implications in longevity.

## Introduction

Nature has been experimenting with the ageing strategies across mammals for more than 200 million years, giving rise to a wide spectrum of lifespans spanning from less than a year (e.g. short-lived shrews) to more than 200 years (e.g. bowhead whales) (George, et al. 1999; Healy, et al. 2014). Typically, there is a positive correlation between body mass and maximum lifespan within mammals. Therefore, the longevity quotient (LQ) was introduced for body mass correction when comparing lifespans across species, which is defined as the ratio of observed longevity to expected longevity for a non-flying mammal of the same body mass (Austad 2010; Austad and Fischer 1991). According to this definition, a few distantly-related lineages, including *Myotis* bats in the order Chiroptera, the naked mole rat (*Heterocephalus glaber*) in Rodentia and the human (*Homo sapiens*) in Primates, exhibit high LQs amongst mammals (Healy, et al. 2014). Their long divergence time implies that extreme longevity has independently evolved multiple times within mammals (Tian, et al. 2017; Wilkinson and Adams 2019; Zhou, et al. 2020).

Over the past two decades, substantial efforts have been made to decipher the molecular mechanisms of mammalian longevity. Genome-wide comparative studies between long-lived and short-lived mammals have revealed a handful of genes and pathways, including nutrient sensing, response to stress, DNA repair and autophagy, that exhibit unique sequence adaptations and age-related expression changes in long-lived species (Gorbunova, et al. 2020; Seluanov, et al. 2018; Zhao, et al. 2021). For example, unique nucleotide changes in *GHR* and *IGF1R* identified in *Myotis brandtii* may alter insulin signalling, possibly contributing to their small body size and long lifespan (Seim, et al. 2013). Emerging evidence has shown that there are only subtle gene expression changes during ageing in *Myotis myotis* and *H. glaber*, and up-regulation of DNA repair and autophagy related genes was observed in these long-lived species, compared to their closely-related short-lived counterparts (Huang, et al. 2020; Huang, et al. 2019; Kim, et al. 2011; MacRae, et al. 2015a). In addition to these findings, a large body of genome-wide studies has also established the associations between candidate genes and longevity, mainly via measuring selective pressure on genes given the phylogeny (Foley, et al. 2018; Tejada-Martinez, et al. 2021; Yu, et al. 2021), detecting differentially expressed genes and proteins between age cohorts or species (Evdokimov, et al. 2018; Fushan, et al. 2015; Huang, et al. 2019; Toren, et al. 2020), and estimating age-related epigenetic changes (Horvath, et al. 2022; Kerepesi, et al. 2022; Wilkinson, et al. 2021). Despite an increase in our understanding of the longevity mechanisms, the link between gene duplication and mammalian lifespan has not been widely investigated.

Gene duplication is considered as a crucial evolutionary mechanism that provides a novel source of genetic variation and thus drives phenotypic innovations (Conrad and Antonarakis 2007; Innan and Kondrashov 2010). Depending on the selective pressure, the extra duplicated copies can be removed to restore the single-copy state, or maintained owing to gene dosage effects or novel mutations that are advantageous for organisms’ survival (e.g. neofunctionalisation and subfunctionalisation) (Birchler and Yang 2022). Several studies have explored gene duplications in the context of cancer resistance, with one of the most remarkable discoveries demonstrating how 20 retrocopies of tumour suppression gene *TP53* contribute to reduced cancer incidence in African elephants (*Loxodonta africana*) (Sulak, et al. 2016). Truncated proteins encoded by 14 of these pseudogenes were suggested to confer cancer resistance by enhancing sensitivity to DNA damage and inducing apoptosis in elephant cells (Sulak, et al. 2016). Likewise, another study identified a higher copy number of two genome maintenance genes, *CEBPG* and *TINF2*, in long-lived naked mole rats compared to humans and mice (MacRae, et al. 2015b). *CEBPG* and *TINF2* are involved in the regulation of DNA repair and telomere protection (Crawford, et al. 2007; Takai, et al. 2011), and their extra gene dosages may allow naked mole rats to better cope with cellular stresses. In addition to production of truncated functional proteins, duplicated pseudogenes can also function as fine-tuning gene expression regulators that modulate the ageing process. For instance, heightened expression of pseudogene *PTENP1* can restore the expression of *PTEN*, a known tumour suppressor, via sponging *PTEN*-targeting microRNAs (e.g. miR-17, miR-20a) in humans cell lines (Li, et al. 2017; Poliseno, et al. 2010). Transgenic mice engineered with an extra *PTEN* copy exhibited enhanced protection from cancer and presented a 16% lifespan extension (Ortega-Molina, et al. 2012). It is speculated that *M. myotis* bats have naturally evolved exceptional longevity by maintaining *PTEN* expression via this mechanism during ageing, compared to short-lived mice (Huang, et al. 2019). These results suggest that investigation of longevity-associated gene duplication can illuminate the molecular basis of longevity evolved in long-lived mammals.

In this study, we investigated the gene duplication events of 287 genes associated with human longevity across genomes of 37 placental mammals. We identified a wealth of genes whose duplication levels exhibit significant correlation with longevity quotients across these species. Noticeably, strong correlation observed for four genes (*CREBBP*, *PIK3R1*, *HELLS*, *FOXM1*) appears to be driven mainly by their high duplication levels in two ageing extremists, the naked mole rat (*H. glaber*) and the greater mouse-eared bat (*M. myotis*). Followed by sequence and expression analyses we further hypothesised that *PIK3R1* gene may have undergone a convergent duplication event in these two long-lived species. Our study identified a few genes whose duplication may underlie the extreme longevity evolved in long-lived mammals, and highlighted the potential role of gene duplication in the evolution of mammalian longevity.

## Results

### Selection of eutherian mammals and human longevity-associated genes used in this study

To ascertain the association between gene duplication and mammalian longevity, the genomes of 37 eutherian species were investigated in this study (Table S1). According to the availability of high-quality genomes these species were selected from eleven orders, representing over 100 million years of evolution. They demonstrate a vast diversity of evolutionary innovations and a wide range of lifespans across placental mammals. The longevity quotients (LQs) range from 0.34 (star-nosed mole, *Condylura cristata*; 55.3 grams) to 5.71 (greater mouse-eared bat, *Myotis myotis*; 28.55 grams) with the median 1.17 ± 1.31 (Table S1). The phylogenetic signal (Pagel’s lambda) of LQs across species was estimated at 0.181 (*P* = 0.572), suggesting that mammalian longevity has evolved independently across taxa. These 37 species were further categorised into four groups based on their LQs (Fig.1a).

A list of 307 human longevity-associated genes from the GenAge database was initially used in this study. These genes have been directly linked to human ageing, and their roles in the ageing process were supported by the findings in model organisms (Tacutu, et al. 2013). After excluding the genes exhibiting paralogous relationships within the dataset (See Materials and Methods, Table S2) and one noncoding gene (*TERC*), 293 genes were retained for the downstream analyses. Using Gene Ontology (GO) enrichment analyses, we noticed that these genes are highly functionally interconnected, and enriched mainly in ageing-related pathways associated with nutrient sensing, response to DNA damage and stress, apoptosis, and phosphorylation and binding (Fig. 1b). Their direct relevance to the ageing process therefore renders these genes as ideal candidates to explore the link between gene duplication and evolution of mammalian longevity.

**Figure 1:**
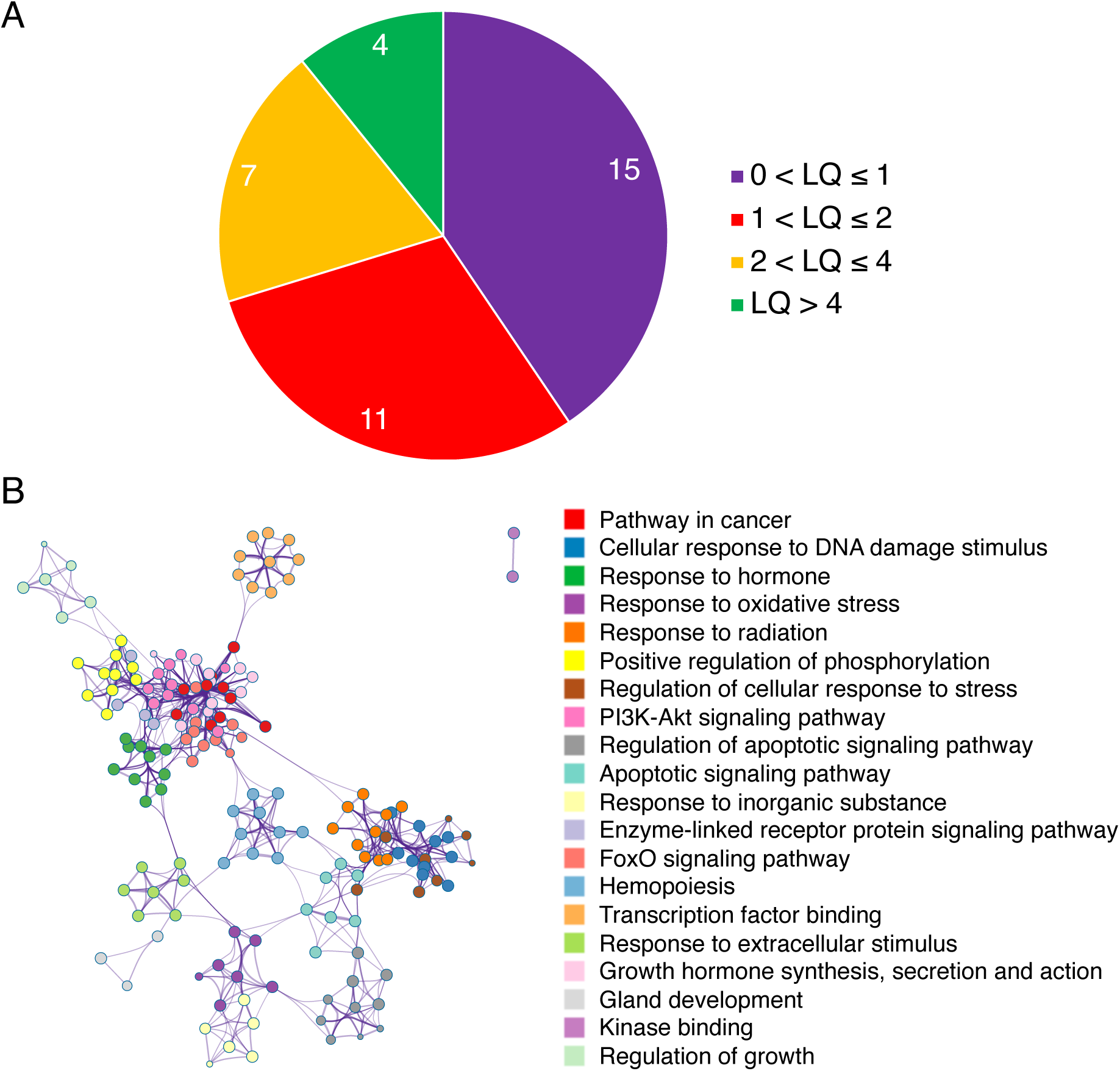
Placental mammals and human longevity-associated genes investigated in this study. (*A*) The distribution of longevity quotients (LQs) across 37 placental mammals. The species are categorised into four groups: 0 < LQ ý 1, 1 < LQ ý 2, 2 < LQ ý 4, and LQ > 4. (B) Gene Ontology (GO) enrichment analysis of 293 human longevity-associated genes. The circles (GO terms) with the same colour indicate that they belong to the same parental term, and the connections between circles indicate that the GO terms share common genes. The size of a circle indicates the number of genes in that GO term.

### Duplication of human longevity-associated in 37 placental mammals

Using discontinuous mega-blast (dc-megablast) we identified the orthologs (RefSeq) of 293 human longevity-associated genes across 37 species and aligned these orthologs against respective genomes to detect their duplication loci (See Materials and Methods). Due to the complexity of gene duplication events (complete or partial duplication) and the disparity of criteria in the degree to which a duplicated fragment is defined as a gene copy (Lallemand, et al. 2020), the duplication level of an ortholog was estimated by the ratio of its cumulative alignment length in a genome to the ortholog length. Using this method, the duplication level of *TP53* was estimated at 20.6 in the African elephant (*L. africana*) genome. This result is consistent with the previous study (Sulak, et al. 2016), suggestive of the reliability of our method. We further removed six genes that have orthologs missing in more than 20% of the species, which resulted in 287 genes retained for further analyses.

We observed that 268 out of 287 genes (93.7%) have the duplication levels, on average, less than 3 across 37 species, with 141 genes (49.1%) having an average of only one copy (original ortholog) (Fig. S1; Table S3). Five genes, including *EEF1A1*, *HMGB1*, *HSPA8*, *YWHAZ* and *HSPD1*, demonstrate high duplication levels (>10) that also exhibit large variations across species (Table S3). We also noticed that a few genes underwent massive expansions in certain species, such as *CHEK2* in *Pan troglodytes*, *IKBKB* in *Echinops telfairi* and *FGFR1* in *Mustela putorius*, whose duplication levels exceed 100. It is important to note that duplicated copies of the genes investigated are mainly truncated gene fragments or retropseudogenes.

Using CAFE analyses, we evaluated the evolutionary trajectories of gene duplications over time. We noticed a considerable difference in the expansion and contraction rates of these 287 genes, with the expansion rate (λ = 0.0052) over 8 times higher than the contraction rate (μ = 0.00056). This is also indicated by the number of genes expanded and contracted on the nodes across the phylogenetic tree (Fig. 2). Interestingly, the ancestral node of Primates, which branches to the species with relatively high LQs (2.39 ± 0.87), exhibits a large number of genes that underwent expansions (+23), while no expansions were observed on the nodes of the clades with a similar divergence time that include species with relatively low LQs, such as Fereuungulata (LQ: 0.95 ± 0.31) and Glires (LQ: 1.32 ± 1.36) (Fig. 2). In particular, 10 and 5 genes were significantly expanded in the two long-lived, divergent species, the naked mole rat (*H. glaber*) and the greater mouse-eared bat (*M. myotis*), respectively (*P* < 0.05; Fig. 2). Notably, the significant expansion of the gene *HELLS* was seen in both species.

**Figure 2:**
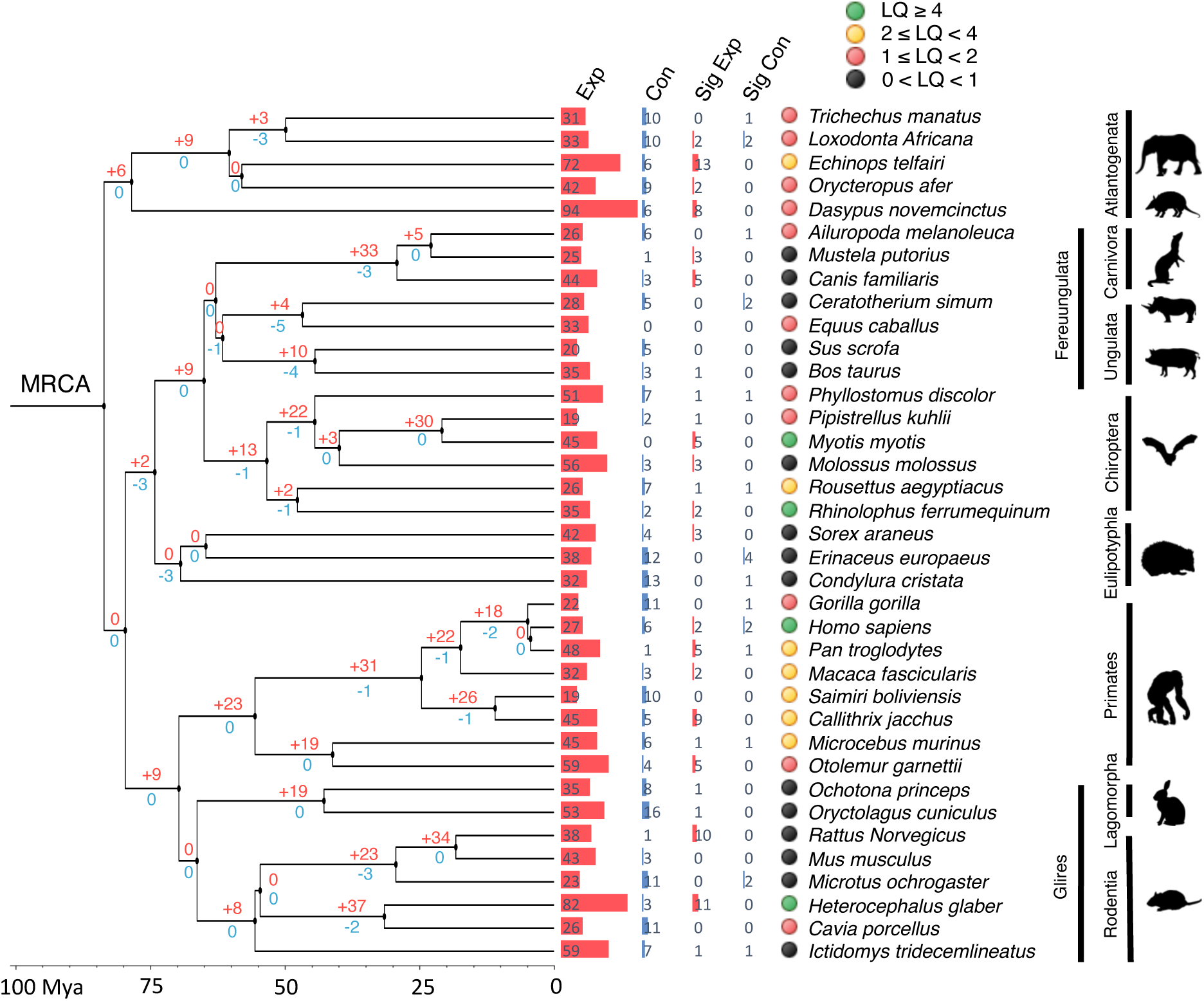
Expansions and contractions of 287 human longevity-associated genes across 37 placental mammals. The time-calibrated phylogenetic tree indicating the relationship amongst 37 mammals was obtained from TimeTree (v5). The values on the phylogenetic tree represent the number of genes expanded (red) and contracted (blue) on each node. The red and blue bars on the tips represent the number of genes expanded and contracted in each species, respectively. ‘Sig Exp’ and ‘Sig Con’ indicate the number of genes significantly expanded and contracted in each species (*P* < 0.05), respectively. The colour code indicates the different LQ groups.

### Phylogenetic correlation analyses between gene duplication and mammalian longevity

Next, we explored the association between gene duplication and mammalian LQ for each gene using phylogenetic correlation analyses (See Materials and Methods). After correcting for phylogeny, 37 genes exhibit significant positive correlation between their duplication levels and LQs (*r* > 0; FDR < 0.05) while only 6 genes show significant negative correlation (*r* < 0; FDR < 0.05) (Fig. 3a; Table S4). The top 3 positively correlated genes are *CREBBP* (*r* = 0.911), *PIK3R1* (*r* = 0.908) and *HELLS* (*r* = 0.897), while the top 3 negatively correlated genes are *HESX1* (*r* = -0.669), *AR* (*r* = -0.601) and *RGN* (*r* = -0.553). However, most of these genes generally have low duplication levels (median: 1.0 ∼ 7.3 across 43 genes; Fig. 3b), and almost all the duplicated copies are truncated pseudogenes. To assess the phylogenetic influence on these analyses, we further performed Spearman’s correlation analyses on the same dataset without phylogeny correction. Only four genes (*AIMF1*, *TFDP1*, *LMNA* and *SOD1*) show significant positive correlation between their duplication levels and mammalian LQs (*r* > 0; FDR < 0.05) (Table S5).

**Figure 3:**
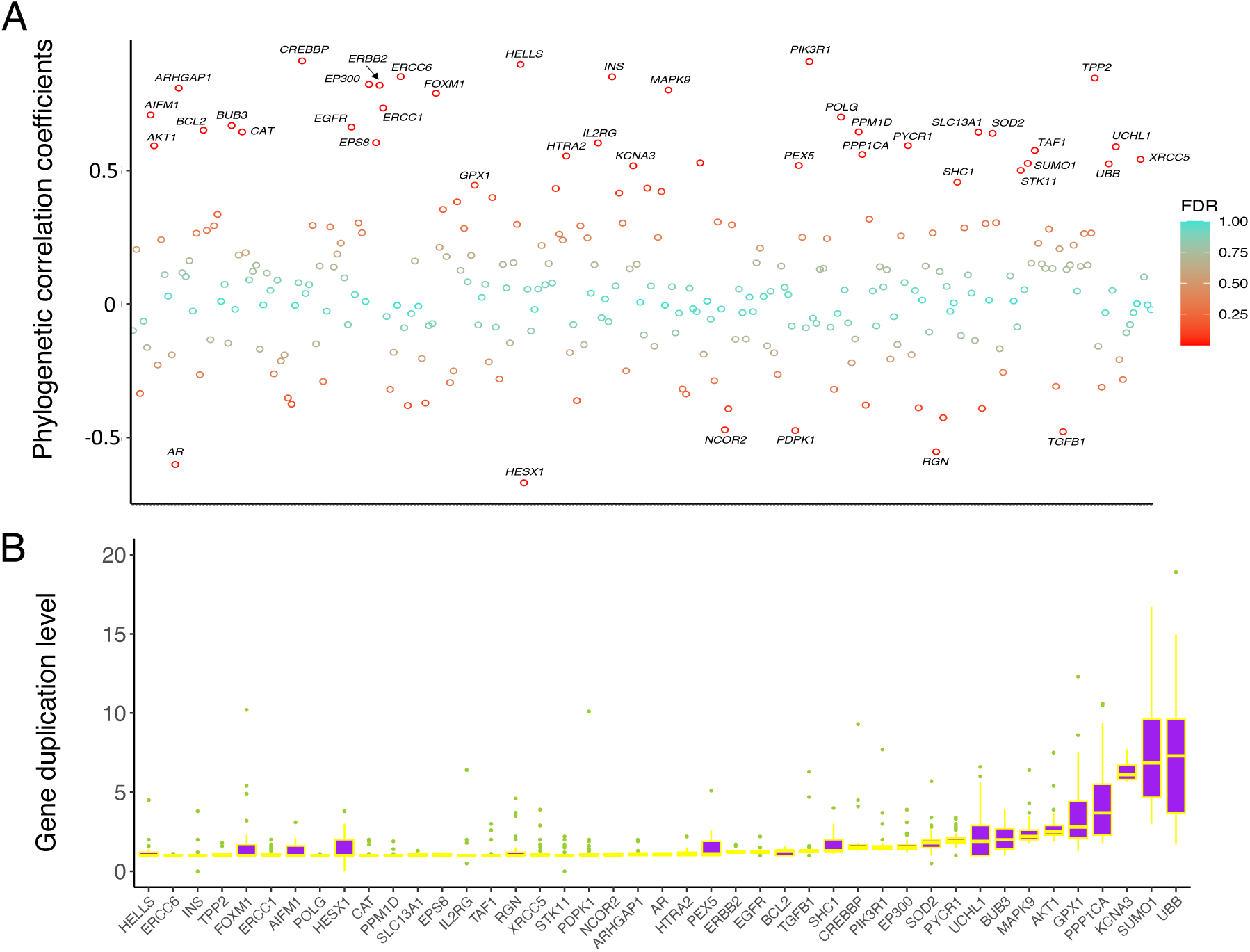
Phylogenetic correlation between duplication levels of human longevity-associated genes and LQ. (*A*) Correlation coefficients of 287 genes after the phylogeny correction. The colour code indicates the significance level (FDR). 43 genes shown on the plot exhibit significant correlation between their duplication levels and mammalian LQs. (*B*) Boxplot showing the duplication levels of 43 significantly correlated genes across 37 species. For each gene, the species with high duplication levels (>20) were not included in the plot.

### Independent duplication of the same longevity-associated genes in long-lived mammals

Out of the 43 genes showing significant correlation between duplication level and mammalian LQ, the strong correlation for four genes (*CREBBP*, *PIK3R1*, *HELLS*, *FOXM1*) results mainly from their high duplication levels in two exceptionally long-lived species, the naked mole rat (*H. glaber*, LQ: 4.59) and the greater mouse-eared bat (*M. myotis*, LQ: 5.71) (Fig. 4). For example, *CREBBP* exhibits a strong positive correlation between its duplication and LQ (*r* = 0.911; FDR = 3.3e^-12^), with *H. glaber* and *M. myotis* having the highest duplication levels (Fig. 4). Likewise, the massive duplication of *PIK3R1* (*r* = 0.908; FDR = 3.3e^-12^) was also observed in these two long-lived species (Fig. 4). Interestingly, we did not find any genes with strong correlation between duplication level and LQ driven by another two long-lived species, the human (*H. sapiens*) or the greater horseshoe bat (*Rhinolophus ferrumequinum*). Next, we investigated the duplication events for *CREBBP*, *PIK3R1*, *HELLS* and *FOXM1* in the *H. glaber* and *M. myotis* genomes. By scanning the duplicated loci on the genomes, for each gene we merged neighbouring loci as a single duplicated copy depending on their corresponding coordinates on the RefSeq (See Materials and Methods). We observed that, for all four genes their duplicated copies are spread across different scaffolds (Fig. 5a). It is noteworthy that, the high duplication level of *HELLS* estimated in the *M. myotis* genome was ascribed to 556 duplications of its last exon (∼85 bp), and this phenomenon might result from transposon activities, such as non-LTR retrotransposition. However, in *H. glaber* the high copy number was attributed to the duplication of different regions of *HELLS*. More excitingly, we found that high copy numbers of *PIK3R1* in both genomes resulted from independent duplications of the similar region of the RefSeq. The coding sequences of *PIK3R1* in both species are 2,175 bp in length. The region (1,000 ∼ 2,100 bp; consisting of last 8 exons) was duplicated 5 times in *H. glaber*, while 13 duplications of the region (1,400 ∼ 2,100 bp; consisting of last 5 exons) were seen in *M. myotis*. Alignments of these pseudogenes demonstrate that they share a common region (∼600 bp) between both species (Fig. 5b). Further genomic analyses show that most of these copies are located in introns, or in intergenic regions in close proximity to protein-coding genes. In the *M. myotis* genome, 3 copies reside in introns, 6 were found in intergenic regions close to 5’ or 3’ UTRs of protein-coding genes (< 30 kb), and 4 are distant (> 30 kb) from protein-coding genes. For *H. glaber*, these numbers are 2, 2 and 1, respectively.

**Figure 4:**
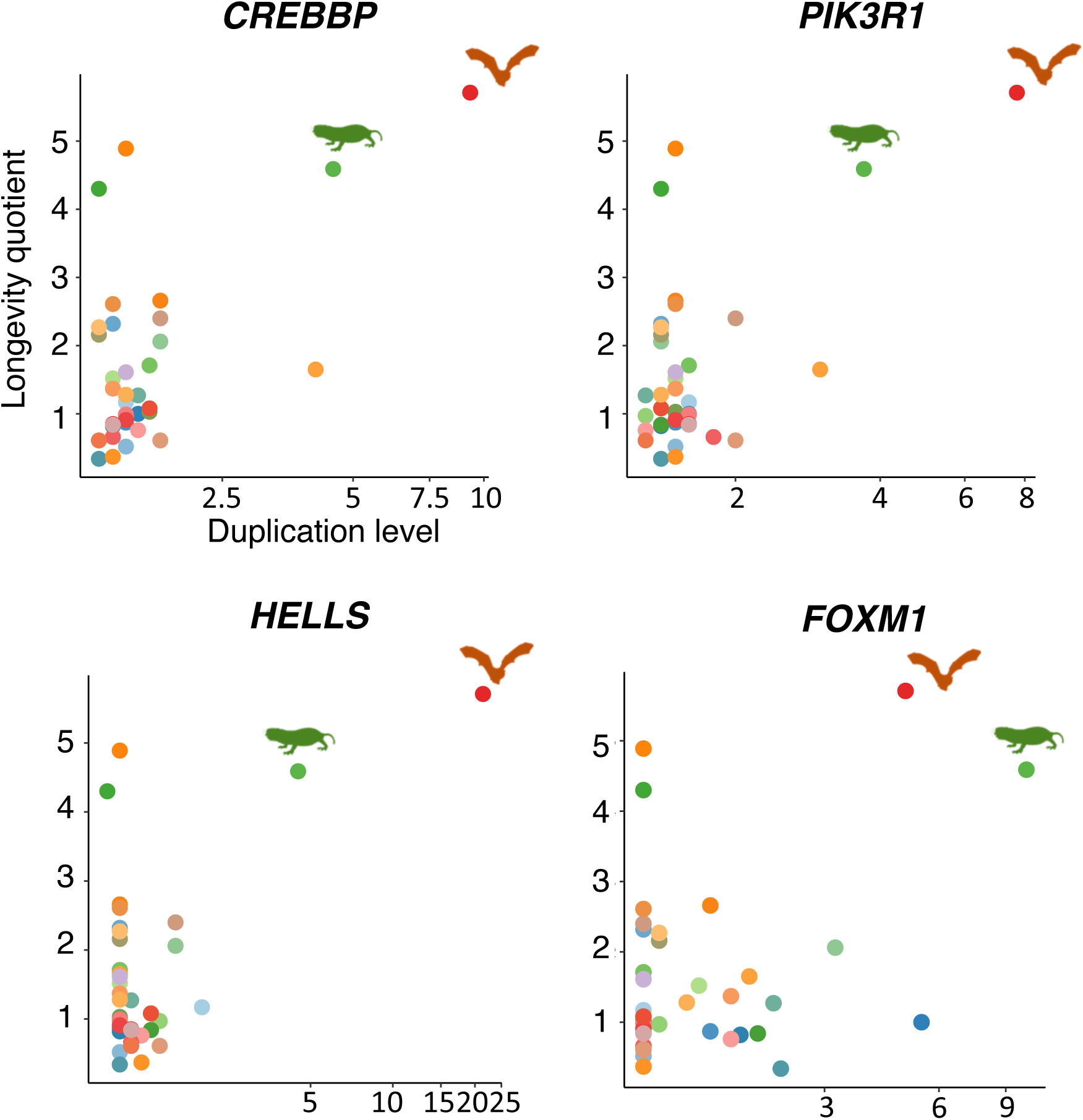
Scatterplots showing the association between gene duplication level and LQ for the genes *CREBBP*, *PIK3R1*, *HELLS* and *FOXM1*. Their strong positive correlation is mainly driven by their high duplication levels in two extremely long-lived species, *H. glaber* and *M. myotis*.

**Figure 5:**
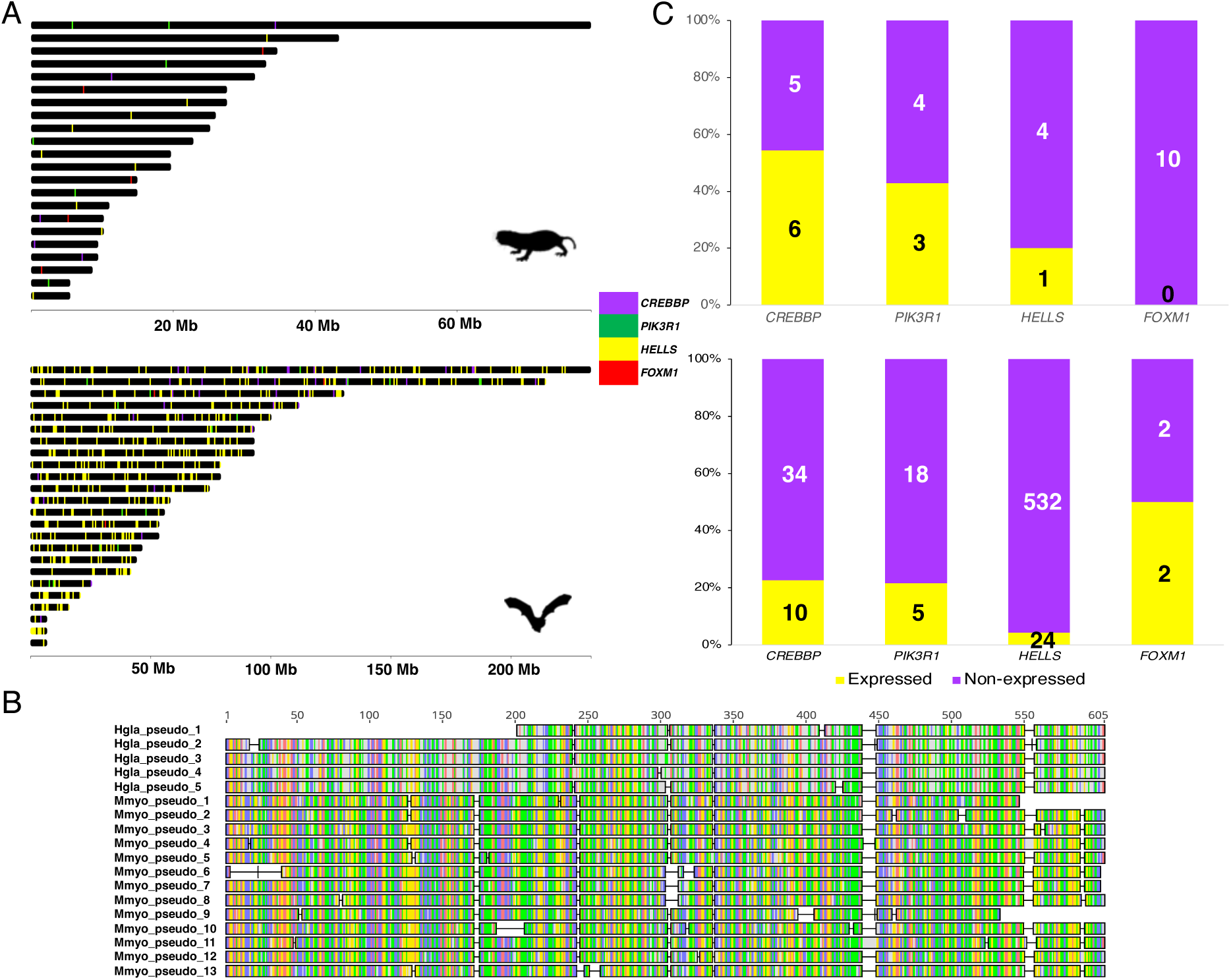
Sequence and expression analyses of four gene candidates in *H. glaber* and *M. myotis*. (*A*) Distribution of duplicated copies of four gene candidates (*CREBBP*, *PIK3R1*, *HELLS*, *FOXM1*) in the *H. glaber* and *M. myotis* genomes. The genome scaffolds shorter than 5 Mb were not included. (*B*) Alignments of *PIK3R1* pseudogenes from *H. glaber* (5) and *M. myotis* (13). Only the ∼600 bp common regions are shown. (*C*) Expression of all the pseudogenes of these four candidates in *H. glaber* and *M. myotis*. The values on the bar plot indicates the number of expressed and non-expressed pseudogenes of each gene in the brain, liver or kidney samples of *H. glaber* and *M. myotis*.

### Expression analyses of duplicated loci of four candidate genes in H. glaber and M. myotis

We further investigated if the duplicated loci of these four genes are expressed, by analysing published brain, kidney and liver RNA-Seq samples from *H. glaber* and *M. myotis*, respectively. We observed that at least one duplicated copy of each gene was expressed in at least one of these three tissues in both species, with the exception of *FOXM1* (Fig. 5c). 54.5% (*CREBBP*), 42.9% (*PIK3R1*), 20% (*HELLS*) and 0% (*FOXM1*) of duplicated copies were considered to be expressed in the naked mole rat, while the percentages are 22.7%, 21.7%, 4.31% and 50% in the greater mouse-eared bat, respectively (Fig. 5c).

## Discussion

### Longevity-associated genes underwent a higher rate of expansions than contractions across the phylogenetic tree

In this study we aimed to ascertain the association between gene duplication and mammalian lifespan. By analysing the evolution of duplication levels of 287 longevity-associated genes across 37 placental genomes, we estimated a much higher rate of expansion (λ = 0.0052) across the phylogenetic tree, compared to their contraction rate (μ = 0.00056) (Fig. 2). This is not surprising because around half (49.1%) of the genes we investigated are single-copy genes across species (Fig. S1). Single-copy genes are functionally important, highly conserved, and are generally expressed at a higher level and in more tissues than non-single-copy genes (De La Torre, et al. 2015; De Smet, et al. 2013). Hence, loss of these single-copy genes likely leads to function loss, thus resulting in detrimental consequences. Meanwhile, unlike the genes with multiple copies, they have been duplication-resistant over the long evolutionary process. This suggests that these single-copy, longevity-associated genes are dosage-sensitive, with their expression levels under strong selective constraints and rigorously regulated.

We observed that a few genes (e.g. *EEF1A1*, *HMGB1*, *HSPA8*) have high and variable duplication levels across species (Table S3; Fig. S1). This phenomenon results from the fact that each of these genes belongs to a large gene family whose members share some conserved domains with each other, and these functional paralogs (family members) underwent further duplication likely through retrotransposition that gave rise to hundreds of thousands of truncated pseudogenes (Troskie, et al. 2021). Meanwhile, high duplication levels of a few genes in certain species (e.g. *CHEK2* in *Pan troglodytes*) were ascribed to large numbers of tandem duplication of short gene fragments. Due to loss of *cis*-regulatory elements and cumulative mutations over time, pseudogenes lose their protein-coding potential and have long been regarded as transcriptionally silent and functionless. However, emerging evidence has challenged this assumption and supports that a proportion of pseudogenes can be translated into truncated proteins or can function as noncoding genes via a number of mechanisms (Cheetham, et al. 2020; Milligan and Lipovich 2014). These include gene silencing by antisense pseudogenes (McCarrey and Riggs 1986), gene silencing by pseudogene-derived siRNA (Tam, et al. 2008), and gene regulation by pseudogenes competing with their parental protein-coding genes for *trans*-acting regulators such as microRNA (Tay, et al. 2014). Nevertheless, owing to unusually high duplication levels, it is unlikely that all these pseudogenes are functional and have dosage effects on physiological consequences. Future experiments to test if they have biological implications at the cellular level are required.

### Evolution of longevity-associated gene duplications likely follows a species-specific manner

If not considering the effect of phylogenetic relatedness, only 4 genes exhibit significant correlation between their duplication levels and LQs (Table S5). After the phylogeny correction, however, 10 times more genes show significant positive or negative correlation (Fig. 3a; Table S4). This result suggests that duplication of a large proportion of longevity-associated genes does not follow phylogenetic patterns and appears to have independent evolutionary trajectories across lineages. Interestingly, it was hypothesised that longevity has independently evolved multiple times during the evolution of mammals (Wilkinson and Adams 2019; Zhou, et al. 2020). Therefore, the strong correlation between gene duplication level and LQ can potentially shed light on the link between gene dosage effect and longevity. However, it is noteworthy that most of these 43 significantly correlated genes have low duplication levels (Fig. 3b), and their duplicated copies are mostly truncated pseudogenes. In addition, due to their species-specific duplication events, it is likely that some of these pseudogenes have not been under natural selection for long enough, so that they may not be fixed in the genomes yet and are unstable (Cardoso-Moreira, et al. 2016). For these reasons we speculated that a large proportion of these duplicated pseudogenes might have no functions, and future investigations are required to test this assumption.

### High duplication levels of the same genes are observed in two long-lived species, the naked mole rat and the greater mouse-eared bat

Four genes (*CREBBP*, *PIK3R1*, *HELLS*, *FOXM1*) exhibit strong positive correlation between their duplication levels and LQs, which is mainly driven by two exceptionally long-lived species, *H. glaber* and *M. myotis* (Fig. 4). *CREBBP* is regarded as a tumour suppressor which plays crucial roles in growth control and homeostasis (Zhang, et al. 2017). Comparative genomic analyses revealed that *CREBBP* is under positive selection in both long- and short-lived mammals suggesting its complex roles in ageing (Yu, et al. 2021). In addition, increased *CREBBP* expression was reported to promote longevity in mice and human populations by activating mitochondrial stress response (Li, et al. 2021). Likewise, *PIK3R1* is also a tumour suppressor and its down-regulation is a prognostic marker in breast cancer (Cizkova, et al. 2013). Interestingly, *HELLS* and *FOXM1* are thought to be involved in cell proliferation, playing essential roles in normal development and survival. However, both genes are considered as oncogenes, whose overexpression promotes metastasis in a variety of cancers (Kim, et al. 2010; Liao, et al. 2018).

Similar to most of the other longevity-associated genes we investigated, all the duplicated copies of *CREBBP*, *PIK3R1*, *HELLS* and *FOXM1* are truncated pseudogenes. Although duplications of the same genes were seen in two extremely long-lived species that are phylogenetically distantly-related, we noted that different genic regions were duplicated in respective species. Noticeably, expression of some of these duplicated loci are supported by RNA-Seq reads sequenced from brain, or liver, or kidney (Fig. 5b), suggesting that these pseudogenes either have potential transcription capability or are the transcriptional by-products of the genes nearby (Harrison, et al. 2005). Nevertheless, even if expressed, these copies may have distinct functions due to their different genic regions derived from the same genes in the respective long-lived species. For example, if acting as microRNA decoys, these expressed pseudogenes will regulate a different set of genes in respective species given the fact that they may possess different microRNA binding sites. Therefore, upon bioinformatic predictions of microRNA binding sites, future cellular experiments, such as luciferase reporter assays, can be used to ascertain if they can regulate expression of different genes by sponging different microRNAs.

### PIK3R1 may have experienced convergent duplications in the long-lived H. glaber and M. myotis genomes

Unlike duplications of *CREBBP*, *HELLS* and *FOXM1*, it is intriguing that the gene *PIK3R1* might have undergone a convergent duplication in the *H. glaber* and *M. myotis* genomes, whereby the similar region of *PIK3R1* was duplicated multiple times in respective species (Fig. 5b). While convergent gene loss that contributes to unique adaptations has been widely reported in animals (Hecker, et al. 2019), convergent gene duplication events that have biological consequences in distantly-related species seem to be rare or overlooked. One striking example comes from the study revealing that the amylase gene (*AMY*) had undergone an expansion in a few distantly-related mammals which consume diet rich in starch (Pajic, et al. 2019). It is therefore important to investigate if the duplicated copies of *PIK3R1* in *H. glaber* and *M. myotis* have biological implications.

To better understand the duplications of the same genic regions of *PIK3R1* in both species, we further investigated its duplication in their closely-related species. We found that the region of *PIK3R1* (1,400 ∼ 2,100 bp) was also duplicated 3 and 18 times in *Pipistrellus kuhlii* (LQ: 1.65) and long-lived *Myotis brandtii* (LQ: 8.23) respectively, but not in any other species investigated. This result indicates that this genic region of *PIK3R1* may have started expansion before the divergence of *Myotis* and *Pipistrellus* bats ∼30 million years ago, and some of these duplications may have been adaptive in their genomes through natural selection over this evolutionary time. However, with the exception of *H. glaber* we didn’t find any duplications of the *PIK3R1* region (1,000 ∼ 2,100 bp) in any other species, including short-lived *Cavia porcellus* which diverged from *H. glaber* ∼40 million years ago. Therefore, it is possible that *PIK3R1* duplication initially occurred in *Myotis* bats and *H. glaber* coincidentally, but acquisition of extra retrocopies may have some evolutionary advantages for their survival so that these duplicated copies have been maintained over time. More closely-related species are needed to better understand the evolutionary trajectories of these duplications.

Owing to lack of introns and the distribution across the genomes, these *PIK3R1* copies are truncated, processed pseudogenes, likely directed by LINE1-mediated retrotransposition (Esnault, et al. 2000). Typically, pseudogenes derived from retrotransposition lack the upstream of the promoters of their parental genes. To obtain transcriptional activity, pseudogenes must be located downstream of a pre-existing promoter in close proximity, or a novel promoter must evolve (Troskie, et al. 2021). By analysing the loci of *PIK3R1* duplications in the genomes of *H. glaber* and *M. myotis*, we noticed that most of them are located either in introns or intergenic regions adjacent to protein-coding genes. Combined with the expression analyses (Fig. 5b), these pseudogenes might share the promoters with their closest protein-coding genes and function as noncoding genes in the post-transcriptional regulation. Nonetheless, one cannot rule out the possibility that pseudogenes distant from protein-coding genes are functional, because we only investigated expressions of these loci in three tissues and other non-coding genes (e.g. long noncoding genes) that possess promoters near these copies are usually poorly annotated in genomes. In the future, it is worth investigating if these truncated pseudogenes can produce functional proteins, analogous to *TP53* duplicated retropseudogenes in African elephants, or if common miRNA binding sites exist on both *PIK3R1* original copy and its pseudogenes, especially the region (∼600 bp) shared by both species. These analyses will enable us to further decipher their potential regulatory mechanisms and functions relevant to mammalian longevity at the cellular level.

## Conclusions

Using comparative genomic approaches, we explored duplications of 287 human longevity-associated genes across 37 placental genomes. We found that nearly half of them are single-copy genes that have been resistant against duplication for a long evolutionary time, suggesting that they are dosage-sensitive and their expression is selectively constrained. We further identified 43 genes whose duplication levels are significantly correlated with mammalian LQs, and most of their duplicated copies are truncated gene fragments or retropseudogenes. Remarkably, we found that strong correlation observed for four genes (*CREBBP*, *PIK3R1*, *HELLS*, *FOXM1*) was driven mainly by their high duplication levels in two exceptionally long-lived species, *H. glaber* and *M. myotis*. In particular, we speculated that the gene *PIK3R1* might have undergone a convergent duplication event, whereby the similar region of its coding sequence was duplicated multiple times in both of *H. glaber* and *M. myotis*. Further sequence and expression analyses indicated the transcription capabilities of some of these pseudogenes. In conclusion, our study revealed several genes whose duplication may contribute to the exceptional longevity evolved in long-lived mammals, potentially providing novel gene candidates for further functional validation of their biological relevance to longevity in the future.

## Materials and Methods

### Longevity-associated genes and species representation

To study the association between protein-coding gene duplication and mammalian longevity, 307 human longevity-associated genes, which are manually curated in the GenAge database (build 20) (Tacutu, et al. 2013), were investigated. The human RefSeq of these 307 genes (the longest transcripts) were obtained from the National Centre for Biotechnology Information (NCBI). Paralogous genes among these 307 genes were identified using a reciprocal BLAST approach (discontinuous mega-blast; dc-megablast) (Altschul, et al. 1990). We considered that two genes are paralogous if more than 50% of their RefSeq were aligned, and we only retained the longest gene for downstream analyses. This is because the presence of paralogs will result in an inaccurate estimation of gene duplication levels in the genome. In addition, we also removed noncoding genes from the list. To understand the biological pathways in which these genes are engaged, we performed gene ontology (GO) enrichment analyses using Metascape (Zhou, et al. 2019).

We investigated duplication levels of human longevity-associated genes in 37 eutherian species across eleven orders. The genome assemblies of these species were downloaded from NCBI and their ecological traits, including body mass (gram), and longevity quotient, were obtained from the previous study (Healy, et al. 2014). To estimate the phylogenetic signal of longevity quotient (LQ) across species, we obtained the time-calibrated mammal tree from TimeTree (v5) (Kumar, et al. 2022) and measured the parameter Pagel’s lambda using *phytools* (Revell 2012). Depending on their LQs, these species were categorised into four groups (0 < LQ <1; 1 χ= LQ < 2; 2 χ= LQ < 4; LQ ≥ 4).

### Estimation of human longevity-associated gene duplication levels in 37 mammalian genomes

To identify the orthologs of human longevity-associated genes in the remaining 36 species, we obtained their RefSeq datasets from NCBI respectively. For each species, we performed sequence alignments with human longevity-associated genes as queries and the RefSeq dataset as the database using dc-megablast (parameters: word_size 11, template_length: 18 template_type coding, *e*-value 1e^-10^). As opposed to blastn and tblastx, dc-megablast enables the identification of distantly-related sequences while avoids inclusions of very short fragments which are considered as false positives. For each gene, the RefSeq with the best hit was considered as orthologous in each species. With these analyses we identified the orthologs of human longevity-associated genes across 36 species.

To estimate the gene duplication level in each species, these orthologs were aligned against the corresponding genome using dc-megablast with the same parameters as mentioned above. Due to the complexity of gene duplication events and the disparity of criteria used to define gene copy number, we introduced a new approach to gauge gene duplication level. The duplication level of an ortholog was defined by the ratio of its cumulative alignment length in the genome to the ortholog length (RefSeq). We removed genes that do not have orthologs in over 20% of the species investigated, and further confirmed their absence in the genomes of respective species using dc-megablast as previously mentioned. In consequence, we obtained a matrix containing the duplication level of each human longevity-associated gene across 37 mammalian genomes (Table S3).

### Expansions and contractions of human longevity-associated genes across the mammal tree

We employed CAFE (v4.2.1) to estimate the expansions and contractions of human longevity-associated genes on the branches given a phylogenetic tree (De Bie, et al. 2006). We firstly rounded gene duplication levels to integers and obtained the time-calibrated phylogenetic tree as aforementioned (Kumar, et al. 2022), which are required as input for CAFE analyses. When running CAFE, error models were applied to correct the estimates of gene duplication levels due to genome assembly or annotation errors, and the gene birth (λ) and death (μ) parameters were separately estimated given the phylogenetic tree and gene duplication levels. A gene with *P*-value < 0.05 indicates that it has a significantly greater expansion or contraction rate of evolution on a certain branch.

### Phylogenetic correlation analyses between gene duplication levels and mammalian longevity quotients

For each gene, phylogenetic correlation analysis between its duplication levels and longevity quotients was performed using the R package *phytools* (Revell 2012). The time-calibrated phylogenetic tree as mentioned above was used to correct correlation tests for the phylogeny. *P*-values were further corrected by false discovery rate (FDR), and a gene with FDR < 0.05 indicates a significant correlation (either positive or negative) between its duplication level and longevity quotient across 37 species. In addition, for the comparison we also performed Spearman’s correlation tests on the same dataset using the R package *cor.test* (Team 2014), without phylogeny correction.

### Sequence and expression analyses of duplication loci of top candidate genes

Next, we focused on four gene candidates (*CREBBP*, *PIK3R1*, *HELLS*, and *FOXM1*) exhibiting strong positive correlation with LQ, which is driven by their high duplication levels in two distantly-related but exceptionally long-lived species (*M. myotis* and *H. glaber*). By scanning the genomic loci in which their duplication occurred, we considered neighbouring loci as a single duplicated copy if they have corresponding continuous RefSeq coordinates within a genomic distance of 200 kb. This is due to the lack of introns in the RefSeqs. The duplicated loci, which have repetitive or overlapping RefSeq coordinates, even if they are adjacent or were produced by retroduplication, are considered as a single duplicated copy. The duplicated loci were visualised in the *M. myotis* and *H. glaber* genomes using the R package *chromoMap* (v0.4.1) (Anand and Rodriguez Lopez 2022). In particular, for the gene *PIK3R1* we also investigated the distance between its duplicated copies and their closest protein-coding genes in *M. myotis* and *H. glaber* respectively, using Bedtools (v2.30.0) (Quinlan and Hall 2010). To further explore *PIK3R1* duplication in closely-related species to *M. myotis*, the genome of another long-lived species (*Myotis brandtii*, LQ: 8.23) was analysed. We further aligned these duplicated copies found in the *M. myotis* and *H. glaber* genomes using Muscle (v5) (Edgar 2004).

To further investigate if these duplicated loci are expressed and thus have potential biological functions, we analysed publicly available RNA-Seq data sequenced from brain, liver and kidney samples of both *M. myotis* and *H. glaber* (Table S6). To do this, for each sample adaptor sequences and low-quality bases (< Q30) were filtered using cutadapt (v3.5) (Martin 2011). Next, clean reads were mapped against the respective genome using HISAT2 (v2.2.1) (Kim, et al. 2019). Alignment files (SAM format) were processed using Samtools (v1.13) (Li, et al. 2009) and Bedtools (v.2.30.0) (Quinlan and Hall 2010), and were further visualised using the genome browser IGV (2.14.1) (Robinson, et al. 2011). A duplicated locus was considered to be expressed if 1) more than 5 reads were mapped onto the junctions between this locus and its flanking regions at both ends; 2) the coverage of this locus by RNA-Seq reads is at least 70%. The open reading frames (ORFs) of these duplicated loci were analysed using Geneious (v11.0.5) (https://www.geneious.com).

## Supplementary Materials

Supplementary data can be found as Figure S1 and Table S1-S6.

## Acknowledgements

We acknowledge the UCD Sonic Performance Computing for the provision of computational resources and support. This work was supported by the Irish Research Council Laureate Bursary Grant (No.74725) and the UCD seed funding (No.68674) to Z.H.

## Data availability

No new data were generated in support of this work. The publicly available genomes and RNA-Seq data used in this study are documented in the Supplemental Tables S1 and S6.

## Figure legends

**Figure S1:** Duplication levels of 287 human longevity-associated genes across 37 mammals. For each gene, the species with high duplication levels (>50) were not included in the plot. The line indicates median duplication level less than 3.

## References

Altschul SF, Gish W, Miller W, Myers EW, Lipman DJ 1990. Basic local alignment search tool. J Mol Biol 215: 403–410.

Anand L, Rodriguez Lopez CM 2022. ChromoMap: an R package for interactive visualization of multi-omics data and annotation of chromosomes. BMC Bioinformatics 23: 33.

Austad SN 2010. Methusaleh’s Zoo: how nature provides us with clues for extending human health span. J Comp Pathol 142 Suppl 1: S10–21.

Austad SN, Fischer KE 1991. Mammalian aging, metabolism, and ecology: evidence from the bats and marsupials. J Gerontol 46: B47–53.

Birchler JA, Yang H 2022. The multiple fates of gene duplications: Deletion, hypofunctionalization, subfunctionalization, neofunctionalization, dosage balance constraints, and neutral variation. Plant Cell 34: 2466–2474.

Cardoso-Moreira M, et al. 2016. Evidence for the fixation of gene duplications by positive selection in Drosophila. Genome Res 26: 787–798.

Cheetham SW, Faulkner GJ, Dinger ME 2020. Overcoming challenges and dogmas to understand the functions of pseudogenes. Nat Rev Genet 21: 191–201.

Cizkova M, et al. 2013. PIK3R1 underexpression is an independent prognostic marker in breast cancer. BMC Cancer 13: 545.

Conrad B, Antonarakis SE 2007. Gene duplication: a drive for phenotypic diversity and cause of human disease. Annu Rev Genomics Hum Genet 8: 17–35.

Crawford EL, et al. 2007. CEBPG regulates ERCC5/XPG expression in human bronchial epithelial cells and this regulation is modified by E2F1/YY1 interactions. Carcinogenesis 28: 2552–2559.

De Bie T, Cristianini N, Demuth JP, Hahn MW 2006. CAFE: a computational tool for the study of gene family evolution. Bioinformatics 22: 1269–1271.

De La Torre AR, Lin YC, Van de Peer Y, Ingvarsson PK 2015. Genome-wide analysis reveals diverged patterns of codon bias, gene expression, and rates of sequence evolution in picea gene families. Genome Biol Evol 7: 1002–1015.

De Smet R, et al. 2013. Convergent gene loss following gene and genome duplications creates single-copy families in flowering plants. Proc Natl Acad Sci U S A 110: 2898–2903.

Edgar RC 2004. MUSCLE: multiple sequence alignment with high accuracy and high throughput. Nucleic Acids Res 32: 1792–1797.

Esnault C, Maestre J, Heidmann T 2000. Human LINE retrotransposons generate processed pseudogenes. Nat Genet 24: 363–367.

Evdokimov A, et al. 2018. Naked mole rat cells display more efficient excision repair than mouse cells. Aging (Albany NY) 10: 1454–1473.

Foley NM, et al. 2018. Growing old, yet staying young: The role of telomeres in bats’ exceptional longevity. Sci Adv 4: eaao0926.

Fushan AA, et al. 2015. Gene expression defines natural changes in mammalian lifespan. Aging Cell 14: 352–365.

George JC, et al. 1999. Age and growth estimates of bowhead whales (*Balaena mysticetus*) via aspartic acid racemization. Can J Zool 77: 571–580.

Gorbunova V, Seluanov A, Kennedy BK 2020. The World Goes Bats: Living Longer and Tolerating Viruses. Cell Metab 32: 31–43.

Harrison PM, Zheng D, Zhang Z, Carriero N, Gerstein M 2005. Transcribed processed pseudogenes in the human genome: an intermediate form of expressed retrosequence lacking protein-coding ability. Nucleic Acids Res 33: 2374–2383.

Healy K, et al. 2014. Ecology and mode-of-life explain lifespan variation in birds and mammals. Proc Biol Sci 281: 20140298.

Hecker N, Sharma V, Hiller M 2019. Convergent gene losses illuminate metabolic and physiological changes in herbivores and carnivores. Proc Natl Acad Sci U S A 116: 3036–3041.

Horvath S, et al. 2022. DNA methylation clocks tick in naked mole rats but queens age more slowly than nonbreeders. Nat Aging 2: 46–59.

Huang Z, Whelan CV, Dechmann D, Teeling EC 2020. Genetic variation between long-lived versus short-lived bats illuminates the molecular signatures of longevity. Aging (Albany NY) 12: 15962–15977.

Huang Z, et al. 2019. Longitudinal comparative transcriptomics reveals unique mechanisms underlying extended healthspan in bats. Nat Ecol Evol 3: 1110–1120.

Innan H, Kondrashov F 2010. The evolution of gene duplications: classifying and distinguishing between models. Nat Rev Genet 11: 97–108.

Kerepesi C, et al. 2022. Epigenetic aging of the demographically non-aging naked mole-rat. Nat Commun 13: 355.

Kim D, Paggi JM, Park C, Bennett C, Salzberg SL 2019. Graph-based genome alignment and genotyping with HISAT2 and HISAT-genotype. Nat Biotechnol 37: 907–915.

Kim EB, et al. 2011. Genome sequencing reveals insights into physiology and longevity of the naked mole rat. Nature 479: 223–227.

Kim HE, Symanowski JT, Samlowski EE, Gonzales J, Ryu B 2010. Quantitative measurement of circulating lymphoid-specific helicase (HELLS) gene transcript: a potential serum biomarker for melanoma metastasis. Pigment Cell Melanoma Res 23: 845–848.

Kumar S, et al. 2022. TimeTree 5: An Expanded Resource for Species Divergence Times. Mol Biol Evol 39.

Lallemand T, Leduc M, Landes C, Rizzon C, Lerat E 2020. An Overview of Duplicated Gene Detection Methods: Why the Duplication Mechanism Has to Be Accounted for in Their Choice. Genes (Basel) 11: 1046.

Li H, et al. 2009. The Sequence Alignment/Map format and SAMtools. Bioinformatics 25: 2078–2079.

Li RK, Gao J, Guo LH, Huang GQ, Luo WH 2017. PTENP1 acts as a ceRNA to regulate PTEN by sponging miR-19b and explores the biological role of PTENP1 in breast cancer. Cancer Gene Ther 24: 309–315.

Li TY, et al. 2021. The transcriptional coactivator CBP/p300 is an evolutionarily conserved node that promotes longevity in response to mitochondrial stress. Nat Aging 1: 165–178.

Liao GB, et al. 2018. Regulation of the master regulator FOXM1 in cancer. Cell Commun Signal 16: 57.

MacRae SL, et al. 2015a. DNA repair in species with extreme lifespan differences. Aging (Albany NY) 7: 1171–1184.

MacRae SL, et al. 2015b. Comparative analysis of genome maintenance genes in naked mole rat, mouse, and human. Aging Cell 14: 288–291.

Martin M 2011. Cutadapt removes adapter sequences from high-throughput sequencing reads. EMBnet. journal 17: pp. 10–12.

McCarrey JR, Riggs AD 1986. Determinator-inhibitor pairs as a mechanism for threshold setting in development: a possible function for pseudogenes. Proc Natl Acad Sci U S A 83: 679–683.

Milligan MJ, Lipovich L 2014. Pseudogene-derived lncRNAs: emerging regulators of gene expression. Front Genet 5: 476.

Ortega-Molina A, et al. 2012. Pten positively regulates brown adipose function, energy expenditure, and longevity. Cell Metab 15: 382–394.

Pajic P, et al. 2019. Independent amylase gene copy number bursts correlate with dietary preferences in mammals. Elife 8: e44628.

Poliseno L, et al. 2010. A coding-independent function of gene and pseudogene mRNAs regulates tumour biology. Nature 465: 1033–1038.

Quinlan AR, Hall IM 2010. BEDTools: a flexible suite of utilities for comparing genomic features. Bioinformatics 26: 841–842.

Revell LJ 2012. phytools: an R package for phylogenetic comparative biology (and other things). Methods in Ecology and Evolution 3: 217–223.

Robinson JT, et al. 2011. Integrative genomics viewer. Nat Biotechnol 29: 24–26.

Seim I, et al. 2013. Genome analysis reveals insights into physiology and longevity of the Brandt’s bat *Myotis brandtii*. Nat Commun 4: 2212.

Seluanov A, Gladyshev VN, Vijg J, Gorbunova V 2018. Mechanisms of cancer resistance in long-lived mammals. Nat Rev Cancer 18: 433–441.

Sulak M, et al. 2016. TP53 copy number expansion is associated with the evolution of increased body size and an enhanced DNA damage response in elephants. Elife 5: e11994.

Tacutu R, et al. 2013. Human Ageing Genomic Resources: integrated databases and tools for the biology and genetics of ageing. Nucleic Acids Res 41: D1027–1033.

Takai KK, Kibe T, Donigian JR, Frescas D, de Lange T 2011. Telomere protection by TPP1/POT1 requires tethering to TIN2. Mol Cell 44: 647–659.

Tam OH, et al. 2008. Pseudogene-derived small interfering RNAs regulate gene expression in mouse oocytes. Nature 453: 534–538.

Tay Y, Rinn J, Pandolfi PP 2014. The multilayered complexity of ceRNA crosstalk and competition. Nature 505: 344–352.

Team RC. 2014. R: A language and environment for statistical computing. R Foundation for Statistical Computing, Vienna, Austria, 2012. In: ISBN 3-900051-07-0.

Tejada-Martinez D, de Magalhaes JP, Opazo JC 2021. Positive selection and gene duplications in tumour suppressor genes reveal clues about how cetaceans resist cancer. Proc Biol Sci 288: 20202592.

Tian X, Seluanov A, Gorbunova V 2017. Molecular Mechanisms Determining Lifespan in Short- and Long-Lived Species. Trends Endocrinol Metab 28: 722–734.

Toren D, et al. 2020. Gray whale transcriptome reveals longevity adaptations associated with DNA repair and ubiquitination. Aging Cell 19: e13158.

Troskie RL, Faulkner GJ, Cheetham SW 2021. Processed pseudogenes: A substrate for evolutionary innovation. Bioessays 43: e2100186.

Wilkinson GS, Adams DM 2019. Recurrent evolution of extreme longevity in bats. Biol Lett 15: 20180860.

Wilkinson GS, et al. 2021. DNA methylation predicts age and provides insight into exceptional longevity of bats. Nat Commun 12: 1615.

Yu Z, et al. 2021. Comparative analyses of aging-related genes in long-lived mammals provide insights into natural longevity. Innovation (Camb) 2: 100108.

Zhang J, et al. 2017. The CREBBP Acetyltransferase Is a Haploinsufficient Tumor Suppressor in B-cell Lymphoma. Cancer Discov 7: 322–337.

Zhao Y, Seluanov A, Gorbunova V 2021. Revelations About Aging and Disease from Unconventional Vertebrate Model Organisms. Annu Rev Genet 55: 135–159.

Zhou X, et al. 2020. Beaver and Naked Mole Rat Genomes Reveal Common Paths to Longevity. Cell Rep 32: 107949.

Zhou Y, et al. 2019. Metascape provides a biologist-oriented resource for the analysis of systems-level datasets. Nat Commun 10: 1523.

