## Supplementary figures and images for "Duplications of human longevity-associated genes across placental mammals"

### Figure S1.pdf

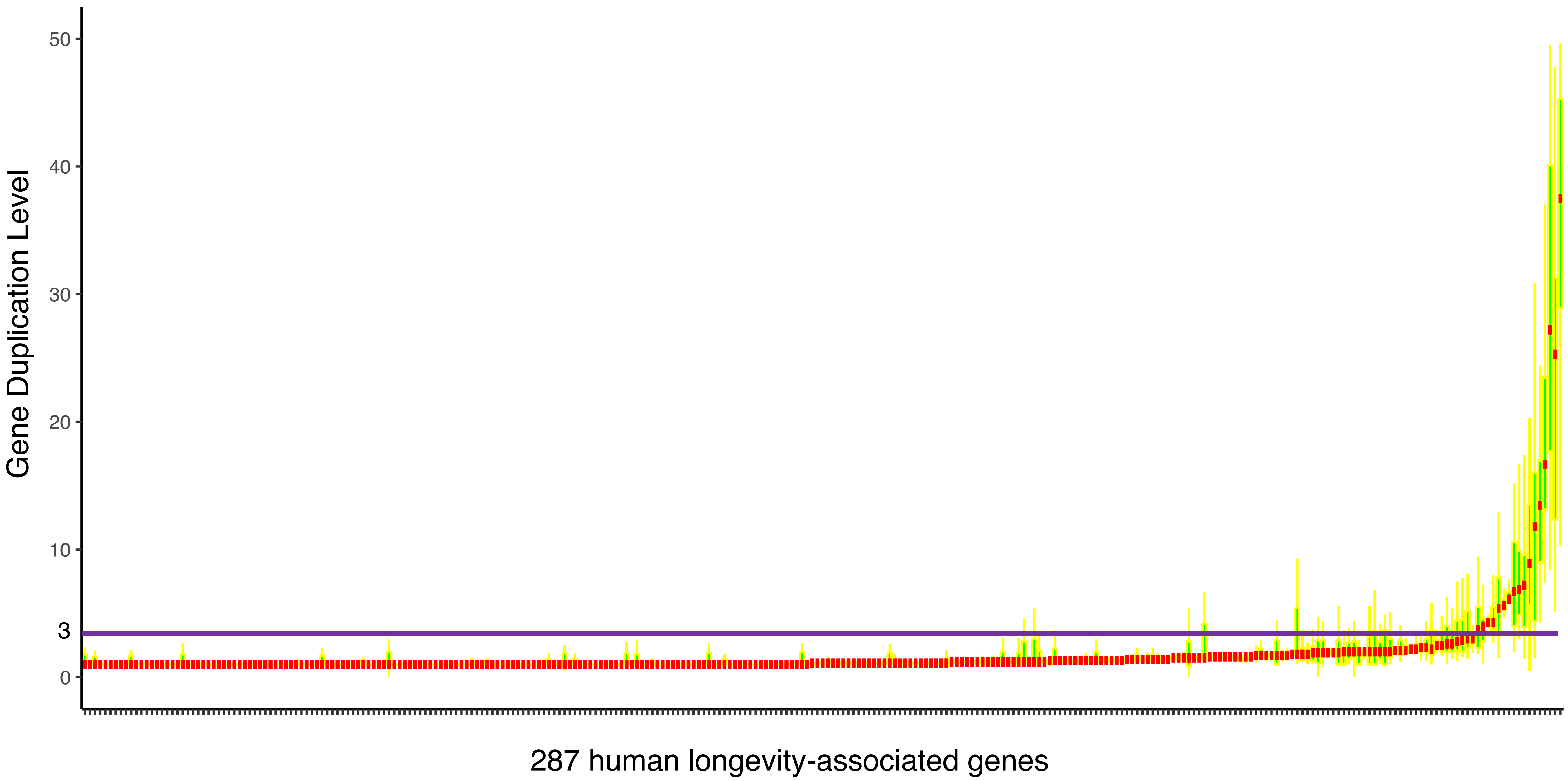
